# Addressing efficacy of everyday hygiene cleansing products in context of sustainable handwashing behaviour in the post-pandemic era

**DOI:** 10.1101/2025.02.13.638075

**Authors:** Sandip B. Pathak, Shafali Arora, Jabir Sayyed, Urmi Trivedi, Lincy Sherin, Nitish Kumar, Harshinie W. Jayasekera, Amitabha Majumdar, Sayandip Mukherjee

## Abstract

Simple hygiene behaviour such as washing hands is key to improving health of individuals and reducing community transmission of communicable diseases such as respiratory and enteric infections. Consistent and relentless messaging by global and local health authorities had resulted in heightened hygiene awareness amongst the public during the COVID-19 pandemic. Unfortunately, many of the hygiene behaviours practiced during the pandemic have proven to be unsustainable in the immediate period following the pandemic. While CDC guidelines suggest washing hands with soap for a minimum of 20 seconds to prevent the spread of germs, a hygiene intervention’s effectiveness must be evaluated in context of the prevalent hygiene behaviour. Here we report our findings from an observational study conducted across India, Pakistan, Saudi Arabia, and United Arab Emirates focusing on handwashing habit of individuals in the post-pandemic period. Across all geographies, the time spent for lathering product on skin (contact time) for a significant majority of individuals recruited for the study was found to be 10 seconds and less. To ensure that marketed hygiene formulations such as liquid cleansers and sanitizers are efficacious in inactivating pathogens under conditions practiced by the majority as a part of their daily hand washing practices, we have investigated the in-vitro antimicrobial efficacy of several hygiene cleansing formulations at 10 seconds of contact time. Our results show that well-formulated cleansing solutions can reduce the input titre of both bacterial and viral pathogens by 99.9% or more, even with brief contact. With the global resurgence of both existing and emerging pathogens in the post-pandemic world, promoting sustainable handwashing practices for infection prevention remains a challenge as it is deeply rooted in the socio-economic fabric of many countries. Therefore, it is crucial to conduct regular studies like this to reassure end-users and combat complacency regarding the composition of everyday hygiene products.

**Highlights:**

- Studies of handwashing behavior in 4 countries reveals significantly short lathering duration
- Antimicrobial efficacy testing of marketed products was conducted against a range of pathogens
- Suitably formulated hygiene products can inactivate pathogens at short durations of contact

## 1. Introduction

In May 2023, the World Health Organization (WHO) declared an end to COVID-19 as a global health emergency. However, ongoing global circulation of multiple sub-lineages of the Omicron variant coupled with recent outbreaks of Mpox virus, H1N1 Influenza virus, Human Metapneumovirus, and Respiratory Syncytial Virus (RSV) across multiple regions of the world continue to underscore the importance of prevention through surveillance and non-pharmaceutical interventions (NPIs) in controlling infectious disease transmission. NPIs are defined as actions, other than medicines and vaccines, that people and communities can take to help slow the spread of communicable illnesses like influenza and COVID-19 (1).

Systematic analysis of data from the Global Burden of Disease survey (1990-2019) revealed stark inequalities between high and low-income countries in terms of relative distribution of non-communicable and communicable diseases (including infectious diseases) (2). While non-communicable diseases constitute greater than 80% of burden in the high-income countries, communicable diseases comprise less than 5% of the total disease burden. In contrast, communicable diseases continue to be the main driver for mortality and morbidity (greater than 60% of total disease burden) in countries across sub-Saharan Africa and South Asia. Considering the five causal groups of communicable diseases (enteric & diarrheal infections, lower respiratory tract infections, HIV, TB, and malaria), India, Nigeria, and Pakistan together account for 48% of disease burden related to enteric infections among children and adolescents, and 44% of lower respiratory tract infections. While the estimates presented above pertain to pre COVID-19, the pandemic has changed the global landscape for communicable diseases and their control and has significantly highlighted the need for preventive interventions such as social distancing and frequent hand--hygiene using soaps, handwashes and alcohol-based sanitizers (3).

There is substantial evidence that shows hand hygiene reduces the spread of several infectious diseases such as gastro-intestinal infections (4–10), respiratory infections (6,11,12), and multiple healthcare associated infections(13,14). The WHO has stated that appropriate hand hygiene (following recommended guidelines) is the most effective action to stop the spread of infections (15). Hand hygiene is a key NPI recommended to prevent the spread of infections in healthcare and community settings alike, with health authorities propagating this habit as a key public health measure for all communicable diseases (16,17).

Hand hygiene recommendations issued by public health organizations, such as the Centers for Disease Control and Prevention (CDC) and the WHO emphasize the critical importance of proper handwashing practices. The CDC advises washing hands with soap and water for a duration of at least 20 seconds, or alternatively, applying alcohol-based hand sanitizer and allowing it to dry for at least 20 seconds (18). Conversely, the WHO’s guidelines for handwashing encompass the entire handwashing procedure in 40 to 60 seconds (19). Although duration of handwashing and hand hygiene practices is suggested to be of 20 seconds or longer (18–20), compliance to this contact time has not been the norm in hand washing behavior(20). Despite the recommendations, studies across different countries and cohorts have shown that people typically spend only a fraction of the recommended time washing their hands, with some studies estimating an average hand washing duration of as little as 6 seconds (20). Other studies have shown that people who got vaccinated during COVID-19 have often changed their preventative behaviors including abandonment of hand sanitation (21).

For the majority, hand washing is a low engagement activity where the decisions regarding whether to wash, the use of product, duration of washing and frequency are left to the individual. This is a challenge that has been long dealt within healthcare (23). Significant progress on hand hygiene compliance has been made over the decades in health care settings that are high contamination and high risk for infection transmission (23–25). Despite stringent training, messaging s and monitoring at workplace, fluctuations in compliance have been reported in healthcare settings including during times of COVID-19 (24). Beyond healthcare settings, widespread external prompts to practice handwashing for prevention, observed during major global outbreaks like the 2009 swine flu and the recent COVID-19 pandemic, have led to intermittent behavior changes, driven by persistent messaging and the perceived threat of infection (22,23). These behavior changes in hand hygiene and other NPIs also resulted in a significant reduction of influenza and other seasonal respiratory infections during the pandemic period (4).

Efforts to promote hand washing for public have primarily focused on educating individuals about the need of using soap and water for prevention of common infections (24). Significant advancements in public health and hygiene standards have been achieved through public and private interventions in these areas (26). Given the well-established role of hand washing towards prevention of diarrheal and respiratory pathogens, especially amidst emerging and re-emerging infections, this simple habit remains crucial and is essential for achieving the United Nation’s Sustainable Development Goal (SDG) target 3.2, which aims to end preventable deaths of newborns and children under 5 years of age, with all countries aiming to reduce neonatal mortality and under 5 mortality (26,27).

In this study, we have employed a cross-sectional study design to investigate the status of adherence to the WHO guideline of 40-60 seconds for handwashing duration, and the CDC guideline of 20 seconds of lathering time. We have also probed the virucidal and bactericidal efficacy of personal hygiene formulations in accordance with the observed handwashing duration.

## 2. Materials and Methods

### 2.1 Handwashing observational study design

A multi-country observational study was conducted with the aim of understanding post pandemic shifts in hand washing behavior using video data collection. To ensure objectivity, the survey was administered by an independent third-party market research agency during the period of Jan 2022 to March 2022. During the study participants were not instructed to use any specific products or given guidelines on how to wash their hands. Their only requirement was to record a video of themselves washing their hands in their household environment.

Data was collected as per ESOMAR28 International Research Code. Participants submitted videos of their general hand washing practice. A total of 901 participants submitted videos. The sample included participants from India, Pakistan, the Kingdom of Saudi Arabia (KSA) and United Arab Emirates within an age range of 18-55 years, including both male and female (Table 1).

**Table 1.**
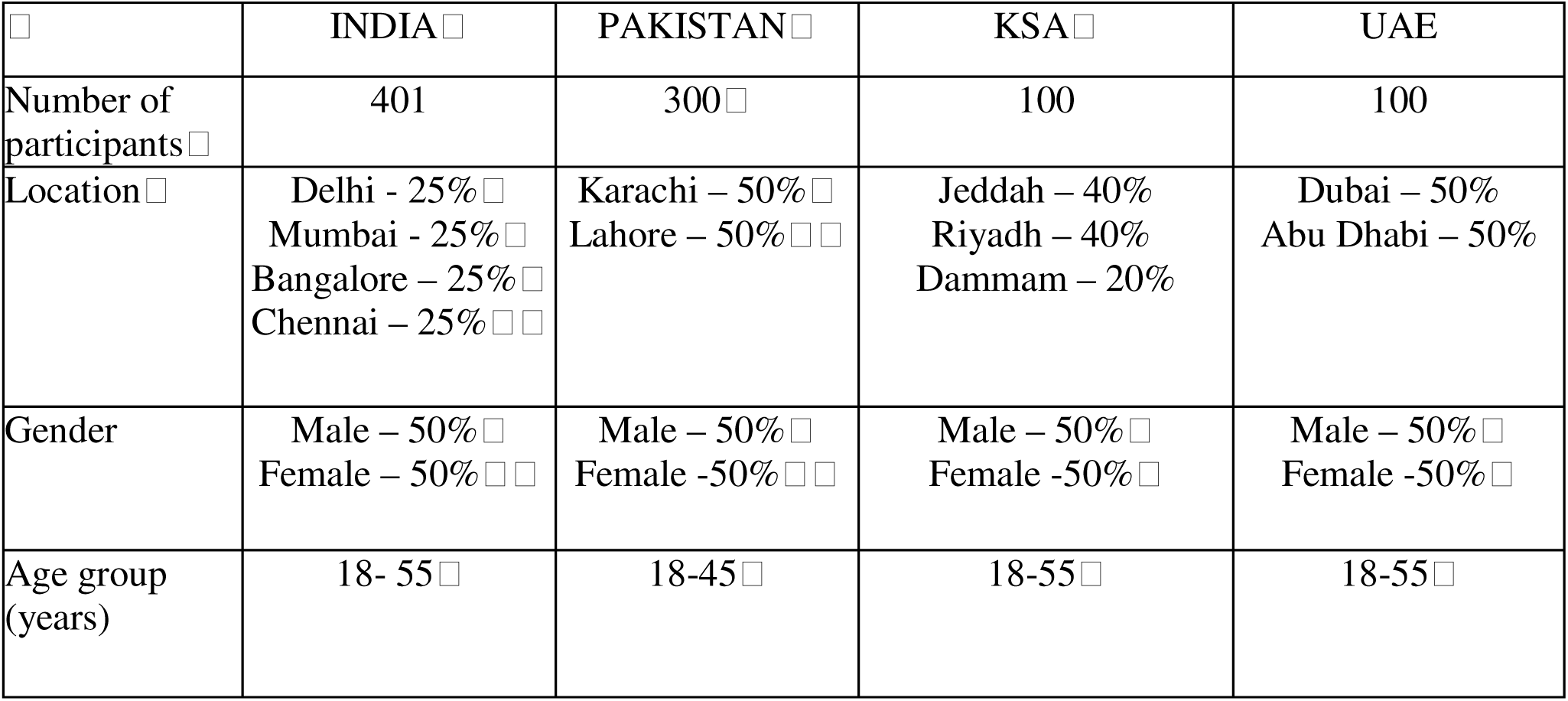
Demographic characteristics of subjects recruited for observational study of handwashing behavior. The table presents the number of participants recruited from four countries: India, Pakistan, the Kingdom of Saudi Arabia (KSA), and the United Arab Emirates (UAE), along with the distribution of participants by location, gender, and age group. Percentages indicate the proportion of participants from each location and gender within each country. Age groups represent the range of participants’ ages in each country.

|  | INDIA | PAKISTAN | KSA | UAE |
| --- | --- | --- | --- | --- |
| Number of participants | 401 | 300 | 100 | 100 |
| Location | Delhi - 25%<br>Mumbai - 25%<br>Bangalore – 25%<br>Chennai – 25% | Karachi – 50%<br>Lahore – 50% | Jeddah – 40%<br>Riyadh – 40%<br>Dammam – 20% | Dubai – 50%<br>Abu Dhabi – 50% |
| Gender | Male – 50%<br>Female – 50% | Male – 50%<br>Female -50% | Male – 50%<br>Female -50% | Male – 50%<br>Female -50% |
| Age group (years) | 18- 55 | 18-45 | 18-55 | 18-55 |

### 2.2 Test products, active ingredients, and test concentrations

The following products were tested for bactericidal and virucidal efficacy: Lifebuoy bodywash; Lifebuoy liquid handwash; Lifebuoy sanitizer gel. The active ingredients in these formulations are as follows: 17% soap (potassium soap) and 3% synthetic detergent (2.1% SLES, 0.75% CAPB) in Lifebuoy bodywash; 8% soap (potassium soap) and synthetic detergent (2.0 % SLES & 2 %CAPB) in Lifebuoy liquid cleanser; 7.8% synthetic detergent (6.3 % SLES, 1.5 % CAPB) and 2% organic acid in Lifebuoy liquid handwash; 70% ethanol in Lifebuoy hand sanitizer gel. All products were manufactured and marketed by Unilever Industries Private Limited. All three ready-to-use commercial products were prepared according to the following test concentrations on the day of the assay: Lifebuoy bodywash and handwash were 50% diluted in distilled water; Lifebuoy hand sanitizer was used undiluted.

### 2.3 Culture, propagation, and enumeration of test bacteria

*E. coli* NCTC 10538, *E. coli* ATCC 10536, *S. enterica* ATCC 14028, *P. aeruginosa* ATCC 15442; *S. aureus* ATCC 6538, and *E. faecalis* ATCC 29212 were used to assess the efficacy of the products previously described against gram-positive and gram-negative bacteria. Culture, propagation and enumeration of test bacteria were conducted in accordance with standard operating procedures for performing in-vitro testing based on the standard method ASTM E2783-22 (26). All the tests were conducted at NABL accredited Bhavan’s Research Centre, Andheri (W), Mumbai, Maharashtra 400058. Approximately 48 hours prior to testing, a sterile tube of Tryptic Soy Broth (TSB) was inoculated from a glycerol stock containing the desired bacteria. The culture was incubated at 35°C± 2°C for 24 hours. Approximately 24 hours prior to testing, the broth culture was inoculated onto the surface of sterile Tryptic Soy Agar (TSA) & incubated at 35°C ± 2°C for approximately 24 hours. A suspension of the challenge microorganism was prepared in 0.9% sterile saline, adjusting the bacterial density to 1.0 × 10^8^ CFU/mL using any technique correlating to aerobic plate count, such as McFarland standard, turbidimetry, or optical density.

### 2.4 Culture, propagation, and enumeration of test viruses

Human respiratory syncytial virus (RSV, ATCC # VR-26) and Influenza A virus H1N1 strain A/PR/8/34 (ATCC # VR-1469) were used to assess the efficacy of hygiene products. RSV were propagated and quantified on commercially obtained Hep-2 cells (ATCC catalogue no. CCL 23) while Influenza A viruses were propagated and quantified using MDCK cells (ATCC catalogue no. CCL 34). Quantification was conducted using standard TCID50 (Tissue Culture Infectious Dose 50) method. Tests using RSV and Influenza virus were carried out at Elements Material Technology Eagan, MN 55121, USA, and at BioScience Laboratory, Bozeman, MT, USA 59718, under biosafety level 2 containment by trained personnel.

### 2.5 Method for determining bacterial reduction efficacy of test products

The evaluation of all test products followed the International standard ASTM E2783-22 (Standard Test Method for Assessment of Antimicrobial Activity for Water Miscible Compounds Using a Time-Kill Procedure)(26). An appropriate aliquot of the challenge suspension (containing respective test organism) was transferred to a vial containing an adequate amount of test product solution and control. The specific bacterial suspension was exposed to the test/product solution and control for a pre-determined exposure/contact time (10 seconds). After the exposure time elapsed, an appropriate aliquot was transferred into neutralizing broth. The sample was then serially diluted and enumerated using the pour plate agar technique and standard microbiological practices. The agar plates were incubated at 35°C ± 2°C for 24-48 hours to yield colony forming units (CFU) per plate. Data is derived to yield CFU/ml and then log transformed for log reduction calculations. All assays included numbers control (water blank) and were validated for neutralization effectiveness. Neutralization solution was found to be effective and nontoxic.

### 2.6 Method for determining virucidal efficacy of test products

All test products were evaluated following ASTM E1052-20 (Standard practice to assess the activity of microbicides against viruses in suspension)(27). Briefly, 1 part virus was added to 9 parts of the diluted/undiluted test product, mixed, and incubated for specified contact time at room temperature. The reaction was then neutralized using a suitable neutralizer before addition to cells. Additionally, the neutralized test solutions were passed through a gel-filtration column (Sephadex LH-20) and the flow-through was collected and serially diluted to add to target cells. Following incubation under cell-culture conditions (37°C and 5% CO2), plates were evaluated for cytopathic effects (CPE) using 50% tissue culture infectious dose. All assays were validated for cytotoxicity control and neutralization control.

## 3. Results

A total of 901 participants (Table 1) completed the study by recording a handwashing episode and sending the video for analysis to an independent third-party research agency. For calculating duration of handwashing, contact time was defined as the total duration of product contact with skin. This included time taken to build up a lather directly with - product interaction as well as the rubbing time of lather on skin until the point it was washed. As shown in table 2, the video analysis revealed the following habits of the cohort recruited for the observational study. In India, 30% of participants washed their hands only with water. They did not use soap bar or a liquid handwashing product and out of the remaining that did use a product, the contact time was 10seconds and less for 50%. Across other geographies, the time spent for lathering product on skin (contact time) for a significant majority is 10 seconds and less with Pakistan at 69%, Saudi Arabia at 79% and 86% reported for United Arab Emirates -as per findings. It is only around 14-26% of participants in all countries included that had a contact time of 11-20seconds between product and hands. A contact time of more than 20 seconds was observed as 10% in India, 5% in Pakistan, 2% in Saudi Arabia with none of the participants employing more than 20 seconds in UAE.

**Table 2.**
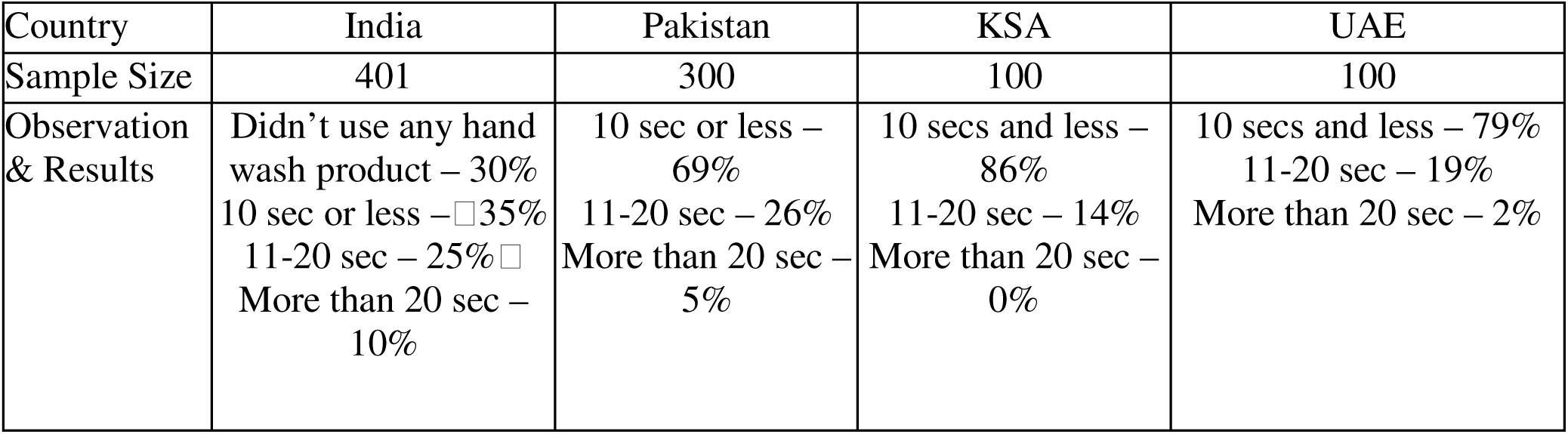
Duration of handwashing with soap based on the observational study of 901 participants across four countries: India, Pakistan, the Kingdom of Saudi Arabia (KSA), and the United Arab Emirates (UAE).

CDC guidelines recommend the use of an alcohol-based hand rub or handwashing with soap and water as a measure to prevent transmission of respiratory viruses including Influenza virus and RSV (28,29). While it is well-established that enveloped viruses are susceptible to the action of surfactants and alcohols present in commercially available hygiene formulations (30), the key parameters that affect the inactivation of viruses include the contact time, the concentration of the disinfecting agent, presence of interfering factors or substances, and the degree of susceptibility of the virus (31). This can be mathematically described by the modified Chick–Watson equation for disinfection 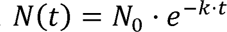 which can also be expressed as 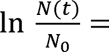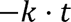 where *N_o_* is the original number of microbes, *N(t)* is the number of viable microbes at time t, t is the contact time, and k is the disinfection rate constant (specific to the microbe) determined by the concentration of disinfectant, susceptibility of the microbe, and environmental factors such as pH and temperature (32). As shown by the equation, a reduction in the contact time t is expected to affect the final log reduction value. Therefore, it is imperative that in view of the changing user habits where the duration of routine handwashing is significantly lesser than the 20 seconds recommended by global health authorities, marketed products are assessed at shorter durations of contact of the microbe with the hygiene formulation to understand their effectiveness in inactivating target microbes. Extending our studies to two common respiratory viruses which have surged globally in the post-COVID years, we observed similar performance at 10 second contact time against both H1N1 and RSV when exposed to the bodywash, handwash and the sanitizer gel (Table 3). The variability in log reduction could be accounted by a combination of factors including differential susceptibility of enveloped viruses to various agents (33), input titer, as well as variable cytotoxicity of the test products to the host cell lines.

**Table 3.** Virucidal efficacy of commercially available hygiene formulations against enveloped viruses at 10 seconds contact duration. Experiments were conducted in triplicate and values indicate average input, output and log reduction. RSV, Respiratory syncytial virus; H1N1, Influenza A virus H1N1 strain.

| Test product & active ingredient | RSV |  |  | H1N1 |  |  |
| --- | --- | --- | --- | --- | --- | --- |
|  | Input titer (Log<sub>10</sub> TCID<sub>50</sub> /mL) | Output titer (Log<sub>10</sub> TCID<sub>50</sub> /mL) | Log<sub>10</sub> TCID<sub>50</sub> reduction | Input titer (Log<sub>10</sub> TCID<sub>50</sub> /mL) | Output titer (Log<sub>10</sub> TCID<sub>50</sub> /mL) | Log<sub>10</sub> TCID<sub>50</sub> reduction |
| Bodywash with soap and synthetic detergent | 8.00 | ≤ 4.75 | ≥ 3.25 | 8.00 | ≤ 4.75 | ≥ 3.25 |
| Handwash with synthetic detergent | 6.25 | ≤ 2.50 | ≥ 3.75 | 7.17 | ≤ 2.50 | ≥ 4.67 |
| Alcohol-based hand sanitizer gel | 6.58 | ≤ 2.50 | ≥ 4.08 | 6.58 | ≤ 2.50 | ≥ 4.08 |

In addition to circulating viruses, both Gram-positive and Gram-negative bacteria are responsible for a wide range of communicable infections transmitted frequently through our hands. Common Gram-positive bacteria like *Staphylococcus aureus* (including MRSA strains) and *Enterococcus faecalis* can cause skin infections, pneumonia, and bloodstream infections. Similarly, Gram-negative bacteria like *Pseudomonas aeruginosa* and *Escherichia coli* can cause urinary tract infections, respiratory infections, and sepsis. Both these classes of bacteria are major causes of community-associated and healthcare-associated infections and are often resistant to multiple antibiotics, making them harder to treat. The role of effective hand hygiene to help prevent the spread of these strains both inside healthcare facilities and in the greater community has been re-emphasized through multiple outbreaks and pandemics (34).To understand if the cleansing formulations tested against enveloped viruses are also effective against Gram-positive and Gram-negative bacteria with a shorter duration of contact, we evaluated the performance of the handwash and the bodywash against a range of representative Gram-positive and Gram-negative bacteria. Both the liquid cleansers achieved a minimum of 3 log_10_ reduction against both Gram-negative and Gram-positive bacteria in 10 seconds of contact time (Table 4). Against specific test bacteria, these cleansers also achieved greater than equal to 4 or 5 log_10_ reduction.

**Table 4.** Bactericidal efficacy of commercially available hygiene formulations against a range of Gram-positive and Gram-negative bacteria. Values indicate mean of three independent experiments along with standard deviation of mean. *These tests were performed with handwash containing 12% synthetic detergent. Remaining tests with handwash were conducted with 7.8% synthetic detergent containing formulations.

| Test products & active ingredient | Log CFU reduction<br>(Mean ± S.D.) | | | | | |
| --- | --- | --- | --- | --- | --- | --- |
|  | *E. coli*<br>10538 | *E. coli*<br>10536 | *E. faecalis*<br>29212 | *P. aeruginosa*<br>15442 | *S. enterica*<br>14028 | *S. aureus*<br>6538 |
| Bodywash with soap and synthetic detergent | Not tested | 4.32 ± 0.08 | Not tested | 5.92 ± 0.01 | Not tested | 3.32 ± 0.18 |
| Handwash with synthetic detergent | 3.05 ± 0.16 | 4.63 ± 0.09* | 5.89 ± 0.02 | 5.91 ± 0.01* | 4.33 ± 0.24* | 5.91 ± 0.08 |

## 4. Discussion

Videography is a powerful tool for collecting behavioral data due to its ability to capture human interactions and behaviors in various contexts. One of the key advantages of videography is its capacity to provide rich and detailed data in naturalistic settings, preserving the authenticity and validity of the observed actions, and eliminating recall bias (35–38). In this study, we utilized videography to observe handwashing practices, including the sequence of actions individuals take during handwashing, the duration of handwashing, whether individuals used soap or not, as well as any deviations from recommended guidelines. We collected and analyzed 901 videos of individuals performing handwashing from India, Pakistan, United Arab Emirates and Saudi Arabia. Our observations were in line with previous reports which have established that despite the positive health benefits, hand hygiene is poorly practiced, - A serial cross-sectional survey assessing COVID-19 knowledge, attitudes, and practices of handwashing in 10 sub-Saharan African countries have shown that handwashing as a COVID-19 prevention strategy notably declined across ten countries through the pandemic, perhaps due to the uptake of other COVID-19 prevention behaviors or due to pandemic fatigue (39). Studies have also demonstrated that decreasing the duration of handwashing and hand rubbing enhances compliance for healthcare workers. For hand rub application, reducing the rub-time from 30 to 15 seconds, and streamlining the technique produced promising outcomes leading to substantially increased compliance in healthcare settings(40). Considering the considerably diminished time of handwashing as observed in our study, it is important to understand if commercially available cleansers are effective enough at reducing microbial load within the short duration of contact of the formulation with the microbes. As mentioned before, handwashing behavior is shifting towards a pre-pandemic state, where contact time between product and skin for the majority is 10 secs or less. Improving long-term adherence to hand hygiene practices, especially in the face of desensitization to health messaging, requires innovative and engaging strategies (41). Therefore, it is important to formulate commonly accessible products for everyday use that provide efficacy to users in congruence to their habit-oriented handwashing behavior. It is also crucial to understand the anti-microbial effectiveness of such formulations in the context of relevant bacteria and viruses which can cause human infections. Given the importance of handwashing in preventing community spread of infections both bacterial and viral in a post pandemic setting, the need for efficacy proven products providing significant pathogen reduction during use could limit the transmission of infection.

## 5. CONCLUSION

Global scale pandemics as seen with COVID-19 are rare and infrequent, but the infection burden of seasonal respiratory viruses, emerging viruses and bacterial pathogens including antibiotic resistant strains remain a constant threat to population. To enhance the effectiveness of handwashing as a daily hygiene practice, especially in reducing the spread of infectious microorganisms, it’s crucial to focus on the efficacy of hand hygiene products within user-relevant contact times. Since public recommendations typically advise using soap or liquid cleanser without specifying the composition, new guidelines which could help address the efficacy gaps among various marketed products would be desirable.

## Acknowledgment

We would like to thank Dr. Samantha Samaras and Dr. Vibhav Sanzgiri for their constant support and encouragement.

